# Neutrophil transit time and localization within the megakaryocyte define morphologically distinct forms of emperipolesis

**DOI:** 10.1101/2021.04.26.441404

**Authors:** Frank Y. Huang, Pierre Cunin, Felix A. Radtke, Ricardo Grieshaber-Bouyer, Peter A. Nigrovic

## Abstract

In emperipolesis, neutrophils transit through megakaryocytes, but it is unknown whether this interaction represents a single type of cell-in-cell interaction or a set of distinct processes. Using an *in vitro* model of murine emperipolesis, we characterized neutrophils entering megakaryocytes using live-cell spinning disk microscopy and electron microscopy. Approximately half of neutrophils exited the megakaryocyte rapidly, typically in 10 minutes or less, displaying ameboid morphology as they passed through the host cell (fast emperipolesis). The remaining neutrophils assumed a sessile morphology, most remaining within the megakaryocyte for at least 60 minutes (slow emperipolesis). These neutrophils typically localized near the megakaryocyte nucleus. By ultrastructural assessment, all internalized neutrophils remained morphologically intact. Most neutrophils resided within emperisomes, but some could be visualized exiting the emperisome into the cell cytoplasm. Neutrophils in the cytoplasm assumed close contact with the platelet-forming demarcation membrane system or with the perinuclear endoplasmic reticulum, as confirmed by immunofluorescence microscopy. Together, these findings reveal that megakaryocyte emperipolesis reflects at least two processes, fast and slow emperipolesis, each with its own characteristic transit time, morphology, and intracellular localization, suggesting distinct functions.

**Key Points:**

- Neutrophil passage through megakaryocytes, termed emperipolesis, diverges into fast and slow forms that differ in transit time, morphology, and intracellular localization
- During emperipolesis, neutrophils can reside in vacuoles (emperisomes) or escape into the cell cytoplasm to assume positions near the megakaryocyte’s demarcation membrane system, endoplasmic reticulum, or nucleus.

## Introduction

Megakaryocytes (MKs) are the largest cells in the bone marrow (50–100 µm) and constitute ∼ 0.05% of marrow cells.^1^ MKs produce platelets by extending long protrusions called proplatelets into sinusoids where shear stress causes platelet release into the circulation.^2,3^ This ability to generate platelets has been extensively studied. However, recent observations have begun to suggest important immune functions.^4,5^ MKs express Toll-like receptors^6-9^ and other immune receptors^10-12^ and produce inflammatory cytokines and chemokines.^13-15^ Early MK progenitors express major histocompatibility complex (MHC) class II.^16^ Mature MKs cross-present antigens to CD8^+^ T cells via MHC I^17^, to CD4^+^ T cells via MHC II^18^, and can exhibit antiviral potency.^19^ In COVID-19 patients, the percentage of MKs in the PBMC fraction is increased and a hyperinflammatory MK subset, enriched in severe COVID-19 patients, constitutes a potential contributor to systemic inflammation.^20^ Thus the functional portfolio of MKs has extended considerably beyond platelet production.

An intriguing functional specialization of MKs is to interact directly with leukocytes, predominantly neutrophils, in a cell-in-cell interaction termed emperipolesis (EP).^21^ Derived from the Greek for “inside round about wandering”, EP was first described in 1956 by Humble et al.^22^ Passage through MKs occurs without apparent harm to either cell.^23,24^ Efficient EP by neutrophils requires active cytoskeletal rearrangement in both the host MK and the transiting neutrophil.^24^ These features clearly distinguish EP from cell-in-cell interactions such as phagocytosis or entosis in which the engulfed cell remains passive and is typically digested.^25^ Under physiological conditions, conventional paraffin sections identify EP in approximately 1– 4% of MKs in mice.^24^ This frequency can more than double with systemic inflammation^24^, chronic blood loss^26^, myelofibrosis^27-29^, myeloproliferative diseases^30^, and gray platelet syndrome.^31-34^

While regularly observed, basic questions regarding the cell biology of EP remain unanswered. We showed previously that neutrophils undergoing EP can fuse transiently with the MK demarcation membrane system (DMS), thereby transferring neutrophil membrane to daughter platelets to enhance platelet production.^24^ Earlier authors had hypothesized that EP may serve as a transmegakaryocytic route for neutrophils in the bone marrow to enter the circulation^26^ or that MKs might provide a “sanctuary” for neutrophils.^35^ Since EP is observed in multiple states of health and disease, the possibility remains that EP could serve several functions.

We hypothesized that, if EP represented a heterogeneous set of processes, then the transit of neutrophils through MKs could exhibit corresponding morphological heterogeneity. We therefore employed immunofluorescence and electron microscopy (EM) to investigate the time course and fate of neutrophils engaged in EP. We demonstrate that EP diverges into fast and slow forms, with multiple distinct intermediate stages, suggesting distinct processes with potentially divergent physiological roles.

## Materials and methods

### Mice

8–12 weeks old WT C57BL/6J mice were purchased from the Jackson Laboratory (#000664) and housed at specific pathogen-free conditions. All animal studies were approved by the Institutional Animal Care and Use Committee of the Brigham and Women’s Hospital.

### Antibodies and reagents

Anti-CD41 APC (MWReg30), anti-CD41 AF488 (MWReg30), anti-Ly6G AF594 (1A8) were from BioLegend. Polyclonal anti-calnexin and FluorSave Reagent were from Sigma-Aldrich. Polyclonal anti-golgin-97, donkey anti-rabbit AF488, DRAQ5, Hoechst 33342, RPMI 1640 with and without phenol red, ACK lysing buffer, paraformaldehyde and glutaraldehyde were from Thermo Fisher.

### Isolation of murine bone marrow cells

Femurs and tibias were flushed with PBS using 22-gauge needles. Cell suspensions were then filtered through 40 µm cell strainers to remove pieces of bone or tissue and centrifuged, followed by lysis of red blood cells using ACK lysing buffer. Bone marrow cells were then washed with PBS and resuspended in complete RPMI medium containing 1% thrombopoietin (TPO medium).

### Isolation of murine megakaryocytes

Hematopoietic progenitor cells were isolated from bone marrow using EasySep™ Mouse Hematopoietic Progenitor Cell Isolation Kit (negative selection) and cultured 1, 2 or 4 days in TPO medium (5×10^6^ cells/ml). Alternatively, bone marrow cells were cultured in TPO medium (10^7^ cells/ml) for 4 days at 37°C, 5% CO_2_. MKs were then enriched using an albumin step gradient.^36^

### Emperipolesis assay

2×10^6^ bone marrow cells and 2×10^4^ MKs were co-cultured in P96 round bottom wells for 12 hours at 37°C, 5% CO_2_.

### Laser scanning confocal microscopy

After 12 hours of co-culture, bone marrow cells and MKs were fixed in PFA 2% for 30 minutes at room temperature. After washing with PBS, cells were resuspended in PBS containing 0.2% saponin and 10% fetal bovine serum (permeabilization buffer) and stained with Hoechst 33342 (5 µg/mL), anti-CD41 AF488 and anti-Ly6G AF594 for 4 hours at RT or overnight at 4°C. In some experiments, cells were stained with Hoechst 33342 (5 µg/mL), anti-CD41 APC, anti-Ly6G AF594 and anti-calnexin or anti-golgin-97 (2.5 µg/mL respectively). Cells were then washed with PBS and resuspended in permeabilization buffer containing donkey anti-rabbit AF488 secondary antibody (10 µg/mL) for 4 hours at RT or overnight at 4°C. After staining, cells were washed and cytospun onto coverslips and mounted on glass microscope slides (Fisher Scientific) using FluorSave Reagent. Images were obtained using a Zeiss LSM 800 with Airyscan attached to a Zeiss Axio Observer Z1 Inverted Microscope using a Plan-Apochromat 63× objective. Zen 2.3 blue edition software was used for image acquisition. Image analysis was performed using ImageJ 1.52p.

### Spinning disk confocal microscopy

Neutrophils and MKs were stained with anti-CD41 AF488 and anti-Ly6G AF594 (1.5 µg/mL respectively) for 1 hour prior to the experiment. DNA was stained with Hoechst 33342 (5 µg/mL) or DRAQ5 (5 µM). Cells were resuspended in TPO medium without red phenol to minimize autofluorescence and plated onto Nunc™ Glass Bottom Dishes (Thermo Fisher) for imaging. Images were obtained using a W1 Yokogawa Spinning Disk Confocal attached to a Nikon Ti inverted microscope with a Plan Fluor 40x/1.3 Oil DIC H/N2 objective and Nikon Elements Acquisition Software AR 5.02 or using a Perkin Elmer Ultraview Vox Spinning Disk Confocal attached to a Nikon Ti inverted microscope with a 60x (1.4NA) objective and Volocity Acquisition Software 6.3. Microscopy chambers were kept at 37°C and 5% CO_*2*_ throughout the experiment. In each experiment, three regions of interest and 10–12 z-stacks were imaged, with approximately 90 seconds between two timepoints. Image analysis was performed using ImageJ 1.52p.

### Tracking of neutrophil migration

The migration of neutrophils through the cytoplasm of MKs was tracked using the ImageJ plugin TrackMate v4.0.1.

### Electron microscopy

After 12 hours of co-culture, bone marrow cells and MKs were washed twice with PBS and fixed in PFA 2% and glutaraldehyde 0.1% for 4 hours at room temperature. Specimens were postfixed in 1% osmium tetroxide and 1.5% potassium ferrocyanide, stained with 1% uranyl acetate, followed by gradual dehydration in 70%, 90%, 100% ethanol and propylene oxide. Specimens were then embedded in Epon. 80 nm sections were imaged using a JEOL 1200EX transmission electron microscope. EM imaging was performed in the Harvard Medical School Electron Microscopy Facility.

### Statistical analysis

Statistical analyses were performed using Graphpad Prism 8 or R. One-way ANOVA and post-hoc Tukey test were performed to compare the frequency of EP across developmental stages. Hartigan’s dip test was performed to determine bimodality in transit time. Unpaired t test was performed to compare neutrophil migration speed between fast and slow EP. All values were displayed as mean ± standard error of the mean. A p-value ≤ 0.05 was considered statistically significant. *p ≤ 0.05, **p ≤ 0.01, ***p ≤ 0.001, ****p ≤ 0.0001.

## Results

### Neutrophil emperipolesis is most efficient in mature megakaryocytes

To study EP, we employed a model wherein murine MKs differentiated from hematopoietic progenitor cells in TPO medium are incubated together with unfractionated murine bone marrow. We performed EP with MKs at different stages in culture, corresponding to different maturational states, grading MK maturity using the extent of DMS.^37^ In each of three experiments, we evaluated EP as performed by 100 MKs at each maturational stage. Mature MKs proved most efficient at EP, displaying neutrophil uptake by approximately 20% of cells in comparison with 0.33% of immature MKs (**Figures 1A–G**). Uptake of > 1 neutrophil was largely restricted to the most mature MKs (**Figures 1H–I**). We therefore employed mature day 4 MK cultures for our studies going forward.

**Figure 1.**
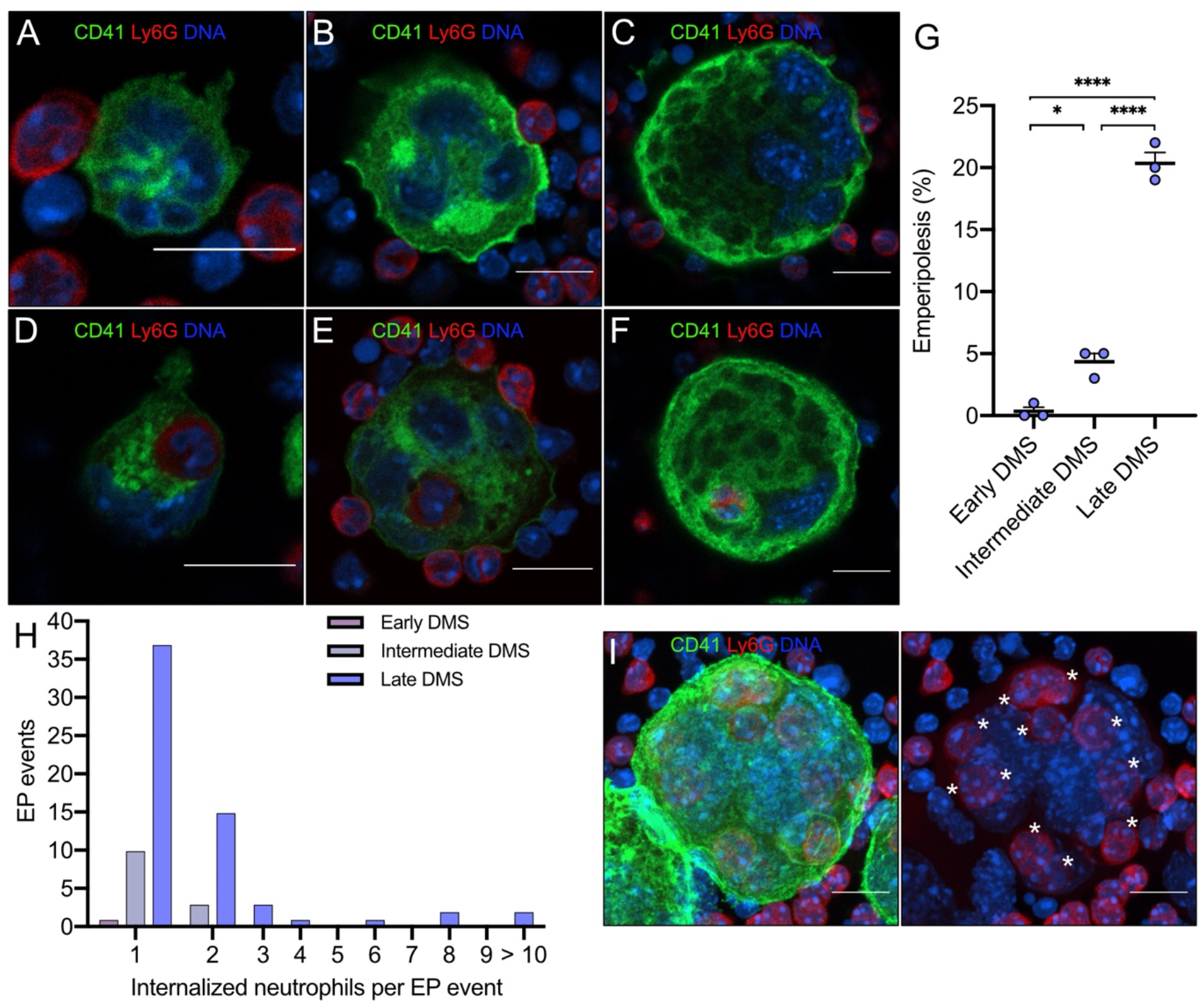
Visualization of EP in MKs of different maturational stages. Hematopoietic progenitor cells were cultured in TPO medium for 1, 2 or 4 days to obtain MKs of different maturation levels. MKs were then co-cultured with bone marrow cells for 12 hours. MKs were stained with anti-CD41 AF488 (green), neutrophils were stained with anti-Ly6G AF594 (red) and DNA was stained with Hoechst 33342 (blue). MK maturity was graded based on the extent of the DMS (early, intermediate and late DMS). Images were obtained using a Zeiss LSM 800 with Airyscan attached to a Zeiss Axio Observer Z1 Inverted Microscope with a Plan-Apochromat 63× objective. Scale bars 10 µm. (A) Immature MK showing DMS beginning to develop between the nuclear lobes and forming connections with the MK surface (early DMS). (B) With increasing maturation, the DMS becomes more prominent and forms thicker connections with the MK surface (intermediate DMS). (C) Mature MK with extensive DMS occupying the majority of the MK cytoplasm (late DMS). (D)–(F) Early, intermediate and late DMS MKs engulfing neutrophils during EP. (G) EP frequency across MK maturational stages. 100 MKs per maturational stage per experiment were counted in 3 independent experiments. (H) The number of engulfed neutrophils per EP event across MK maturational stages. Pooled data from three independent experiments (n = 300 MKs per maturational stage, EP events: early DMS MKs: 1, intermediate DMS MKs: 13 and late DMS MKs: 61). (I) Z-projection of mature MK (late DMS) containing 11 neutrophils (asterisks).

### Neutrophil transit time through megakaryocytes is bimodally distributed: fast and slow emperipolesis

To understand whether neutrophil transit through MKs represents a uniform or heterogeneous process, we employed spinning disk confocal microscopy. MKs were cultured together with whole bone marrow cells and visualized over 90 minutes, obtaining images every 90 seconds and acquiring 10–12 z-stacks per MK to distinguish internalized from superimposed neutrophils. We analyzed 28 EP events. Half (14/28) of neutrophils completed passage through MKs in 30 minutes or less (**Figures 2A–B; Supplementary Movie 1**). Of these “fast emperipolesis” events, most (11/14; 79%) lasted less than 10 minutes, with some episodes of EP occurring in as little as 3 minutes. By contrast, other neutrophils resided within MKs for at least 50 minutes (13/28; 46%), with most (12/13; 92%) remaining for > 60 minutes (“slow emperipolesis”) (**Figures 2A and C; Supplementary Movie 2**). Hartigan’s dip test^38^ confirmed a bimodal distribution of transit times (p = 7.89 x 10^−7^). Interestingly, fast and slow EP could be observed simultaneously in the same MK (**Figure 3A**). These findings show that experimental EP encompasses fast and slow forms with distinct time courses.

**Figure 2.**
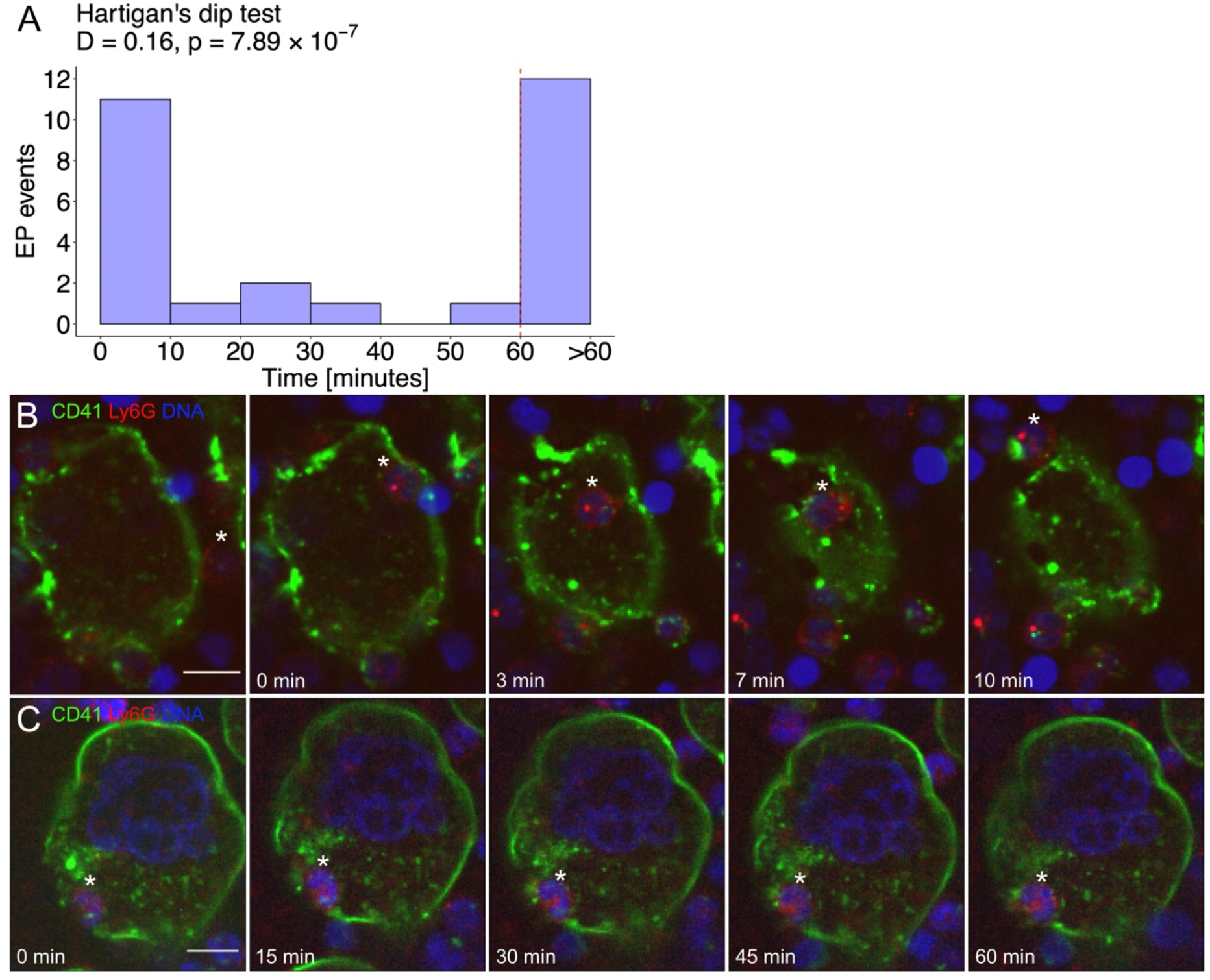
Neutrophil transit time through MKs is bimodally distributed: fast and slow EP. Mature MKs were stained with anti-CD41 AF488 (green) and co-incubated with fresh bone marrow cells stained with anti-Ly6G AF594 (red). DNA was stained with DRAQ5 (blue). (A) Histogram depicting the duration of neutrophil transit through MKs of 28 EP events reveals a bimodal distribution with peaks between 0–10 minutes (fast EP) and > 60 minutes (slow EP). Results pooled from 5 independent experiments. Bimodality was confirmed by Hartigan’s dip test (D = 0.16, p = 7.89 x 10^−7^). (B)–(C) Images were obtained using a W1 Yokogawa Spinning Disk Confocal attached to a Nikon Ti inverted microscope with a Plan Fluor 40x/1.3 Oil DIC H/N2 objective. Scale bars 10 µm. (B) Representative image sequence of fast EP. The neutrophil (asterisk) enters the MK on the right side, migrates through the MK cytoplasm and egresses on the opposite side within 10 minutes. (C) Representative image sequence of slow EP. The neutrophil (asterisk) is already inside the MK at the beginning of the image acquisition and remains inside for at least 60 minutes showing no migration inside the MK.

**Figure 3.**
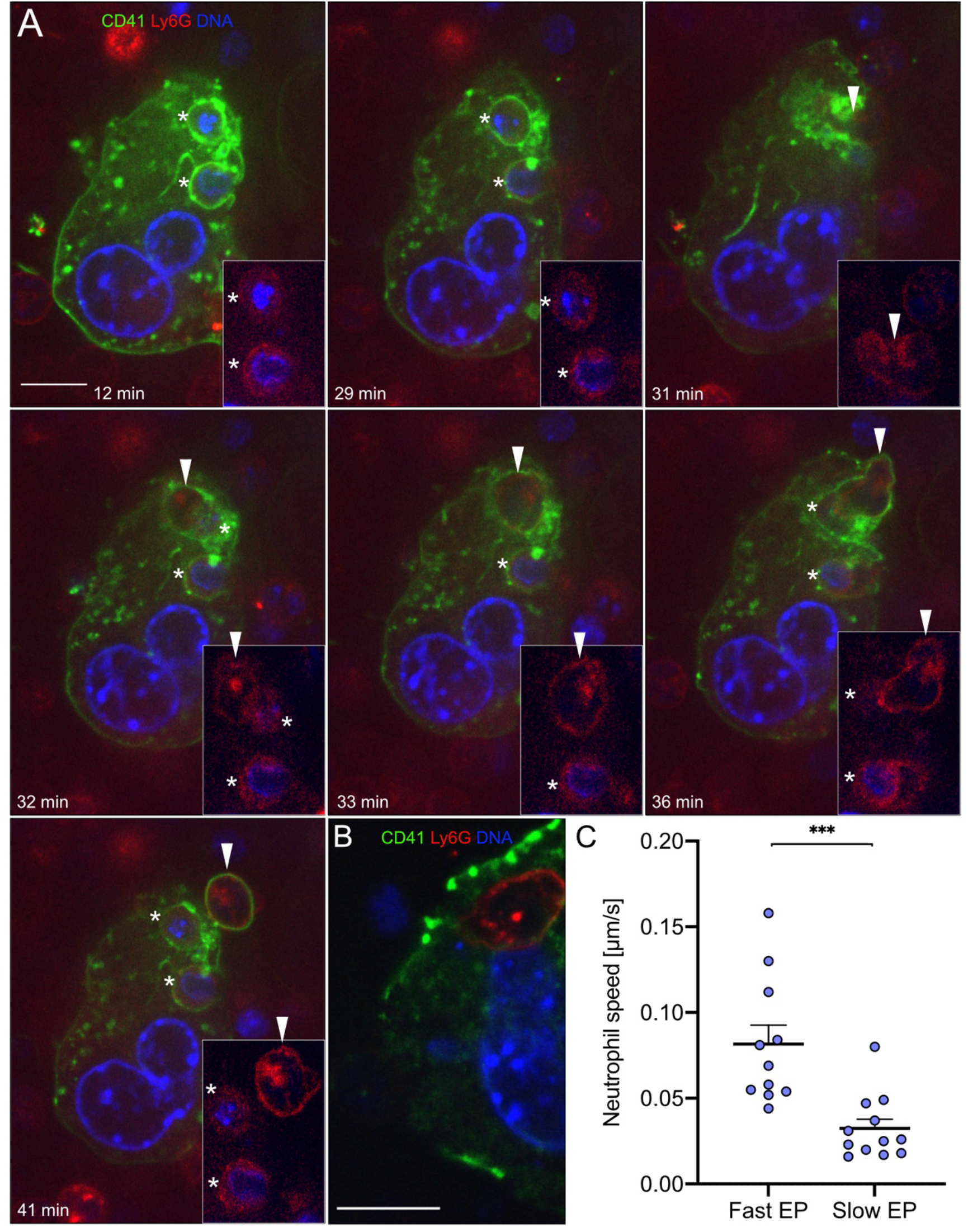
Distinct morphology of fast and slow EP within a single MK. Mature MKs were stained with anti-CD41 AF488 (green) and co-incubated with fresh bone marrow cells stained with anti-Ly6G AF594 (red). DNA was stained with Hoechst 33342 (blue). (A)–(B) Images were obtained using a Perkin Elmer Ultraview Vox Spinning Disk Confocal attached to a Nikon Ti inverted microscope with a 60x (1.4NA) objective. Scale bars 10 µm. (A) An MK showing fast and slow EP simultaneously to illustrate morphological differences of both forms. Two neutrophils undergoing slow EP (asterisks) assume a sessile state. A third neutrophil (arrowhead) enters the MK after 31 minutes and extends dynamic membrane protrusions to propel itself through the MK cytoplasm, exiting within few minutes. (B) Representative image of a slow EP neutrophil residing near the MK nucleus. (C) The passage of neutrophils undergoing fast EP (< 10 minutes) and slow EP (> 60 minutes) through MKs was tracked using the ImageJ plugin TrackMate to determine the mean speed of migration.

### Distinct morphology of fast and slow emperipolesis

We characterized the morphological features of fast and slow EP. Neutrophils undergoing fast EP rapidly transited through MKs by extending dynamic membrane protrusions, appearing to propel themselves through the host cell (**Figure 3A; Supplementary Movie 3**). By contrast, in slow EP, resident neutrophils remained in a single location within the MK for the entire duration, without any signs of mobility (**Figure 3A; Supplementary Movie 3**). These long-term resident neutrophils localized adjacent to the MK nucleus in 42% of events (5/12), in some cases showing close approximation to the nucleus (**Figure 3B**). We then tracked neutrophil movement within MKs for fast and slow EP. Neutrophils undergoing fast EP showed a 2.5-fold higher mean migration speed inside MKs compared to neutrophils undergoing slow EP (**Figure 3C**). These results confirm our observation that slow EP neutrophils remain largely sessile. Fast and slow EP are thus morphologically as well as chronologically distinct.

### Characterization of the ultrastructural features of emperipolesis by electron microscopy

Prior studies have shown that neutrophils engaged in EP may reside either in MK vacuoles, termed emperisomes, or directly within the MK cytoplasm.^24^ We sought to better understand the relationship between these compartments in EP using transmission EM. We co-cultured murine MKs and bone marrow cells as above and processed them for EM after 12 hours of culture, analyzing 45 EP events across five independent experiments.

All MKs and neutrophils remained morphologically intact, without membrane blebbing, nuclear fragmentation, or other evidence of apoptosis. Residence within an emperisome was the most common location for internalized neutrophils (18 of 45 events, 40%), yet the interaction between neutrophil and emperisome was heterogeneous. In some cases, the emperisome membrane was smooth and separated from the neutrophil by a large pericellular space (**Figure 4A_I_**). Alternately, the emperisome membrane could exhibit small protrusions extending towards the engulfed neutrophil (**Figure 4A_II_**). Finally, some emperisomes were tightly wrapped around the neutrophil, with zipper-like approximation of neutrophil and emperisome membranes (**Figure 4A_III_**). Intriguingly, some neutrophils appeared within a vacuolar space but in contact with the cytoplasmic DMS without an interposed MK membrane, suggesting an egress event mediated by reshaping or dissolution of the emperisome (**Figure 4B**_**I**_). Others were entirely surrounded by DMS (**Figure 4B_II_**). Importantly, some neutrophils could be visualized transiting directly from an emperisome into the MK cytoplasm, far from the DMS (**Figure 4C_I_**), or fully resident within the MK cytoplasm without any interposed MK membrane (**Figure 4C_II_**). The frequencies of these different EP stages are shown in **Figure 4D**. Of note, despite the proximity of the neutrophil to the MK nucleus in some instances, our EM studies identified no examples of direct membrane-membrane contact. While the fixed nature of EM images precludes assignment of these stages to fast or slow EP, these images confirm the highly varied interaction between an internalized neutrophil and its MK, suggesting further that penetration through the emperisome membrane represents the most common mechanism of neutrophil egress into the MK cytoplasm.

**Figure 4.**
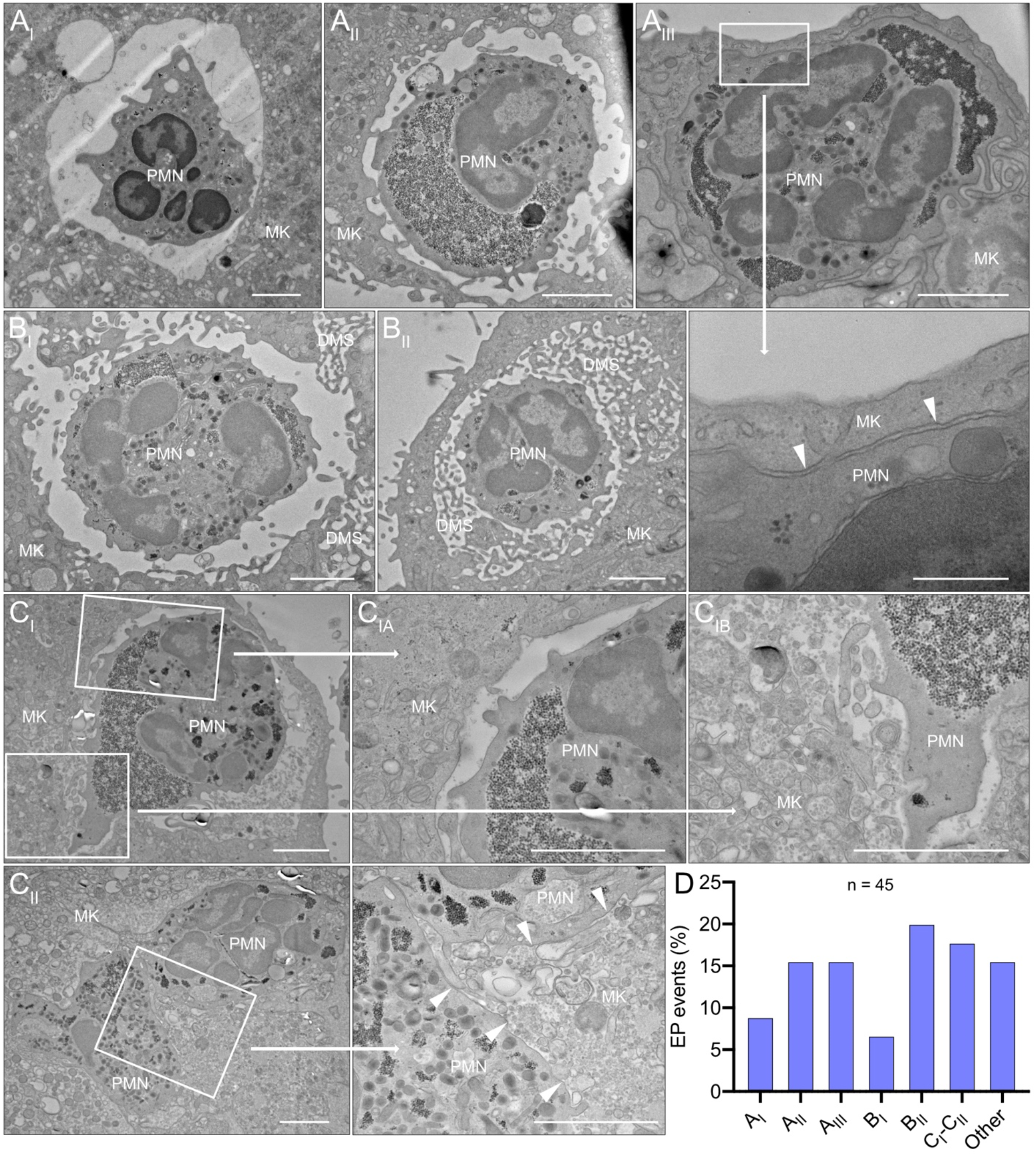
Characterization of the ultrastructural features of EP by electron microscopy. Mature MKs incubated with bone marrow cells for 12 hours were fixed and processed for transmission EM. 45 EP events were observed from five experiments. (A_I_)–(C_II_) Transmission EM images of EP. Scale bars 2 µm or (for magnification of A_III_) 500 nm. (A_I_) Large round emperisome with a smooth vacuolar membrane surrounding a neutrophil (PMN). (A_II_) The emperisome extends membrane protrusions toward the engulfed neutrophil. (A_III_) The emperisome tightly wraps around the engulfed neutrophil. Magnification shows close membrane approximation between neutrophil and emperisome membranes (arrowheads). (B_I_) Internalized neutrophil partly covered by the emperisome and partly exposed to the DMS of the MK. (B_II_) Neutrophil residing within the cavities of the DMS. (C_I_) Internalized neutrophil partly covered by the emperisome (C_Ia_) and partly exposed to organelles of the MK cytoplasm (C_Ib_). (C_II_) Two neutrophils fully reside inside the MK cytoplasm. Only the neutrophil membranes remain visible (arrowheads). (D) Frequency of the previously described EP stages (n = 45).

### Neutrophils interact with the megakaryocyte endoplasmic reticulum as well as the DMS

The DMS is easily recognized by its dilated appearance (**Figure 5A**, bottom left), but not all interactions between cytoplasmic neutrophils and MK organelles were with the DMS. Identifying membranes as belonging to the endoplasmic reticulum (ER) or the Golgi apparatus can be difficult by EM. 7 of 45 neutrophils (16%) were surrounded by membranes that we could not unambiguously assign to one of these structures (**Figures 4D and 5A**). Given our observation that neutrophils undergoing slow EP frequently reside near the MK nucleus, we hypothesized that they might interact with the perinuclear ER. Immunofluorescence staining confirmed that the perinuclear ER sometimes surrounded intracytoplasmic neutrophils (**Figures 5B–C**), enclosing neutrophils between ER and nucleus. This location is distinct from that of the DMS, as reflected in the inverse distribution of the calnexin^+^ ER and the CD41^+^ DMS (**Figures 5B–C**). The Golgi apparatus, another intracellular membrane network marked by golgin-97, did not colocalize with internalized neutrophils (**Figure 5D**). These data further confirm the diversity of intramegakaryocytic localization by neutrophils in EP, supporting the heterogeneity of this process.

**Figure 5.**
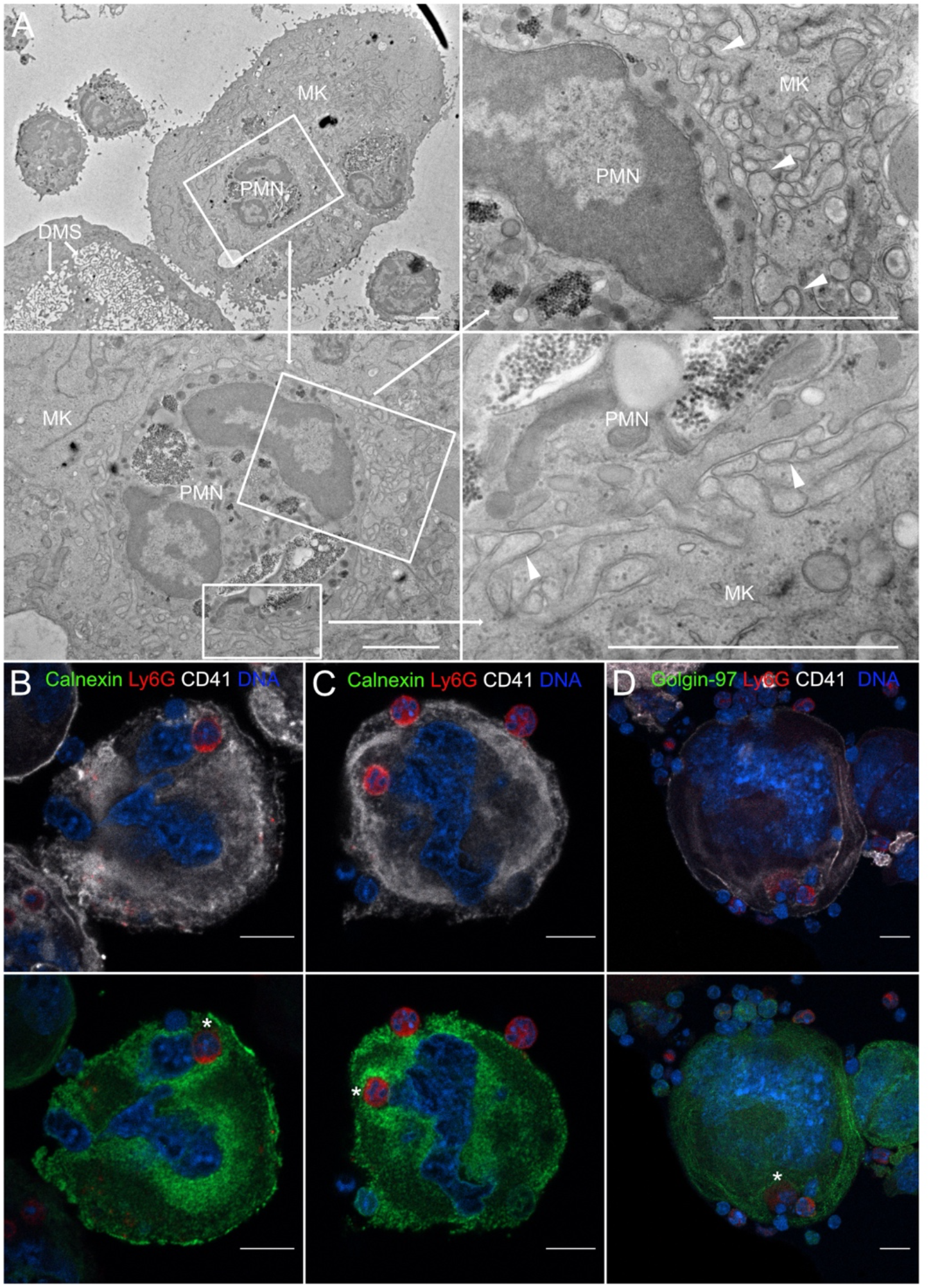
Neutrophils interact with the MK endoplasmic reticulum as well as the DMS. MKs were allowed to engage in EP for 12 hours followed by processing for EM or laser scanning confocal microscopy. (A) Transmission EM images of EP. The internalized neutrophil is surrounded by a membrane network (arrowheads) that does not resemble the DMS. Scale bars 2 µm. (B)–(D) Cells were stained with anti-CD41 APC (white), anti-Ly6G AF594 (red), anti-calnexin or anti-golgin-97 (green) and Hoechst 33342 (blue). Images were obtained using a Zeiss LSM 800 with Airyscan attached to a Zeiss Axio Observer Z1 Inverted Microscope with a Plan-Apochromat 63× objective. Scale bars 10 µm. (B) and (C) The perinuclear portion of the MK endoplasmic reticulum (ER) surrounds the internalized neutrophils (asterisks). The neutrophils are localized between the MK nucleus and ER. Note the inverse distribution of the DMS and ER. (D) Internalized neutrophils (asterisk) did not colocalize with the Golgi apparatus of MKs.

## Discussion

Emperipolesis is a cell-in-cell interaction at the interface of hemostasis and immunity. Neutrophils pass through MKs without disrupting the integrity of either cell, penetrating into the MK cytoplasm in at least some cases. Other granulocytes, lymphocytes, erythrocytes or monocytes are occasionally observed inside MKs, but neutrophils predominate and are observed more frequently than any other lineage even after adjusting for their abundance in the bone marrow.^24,39,40^

Despite the frequency of EP, it remains unknown why neutrophils enter MKs, whether different forms of EP exist, and what roles this interaction plays in health and disease. Understanding the cell biology of EP will be key to answering these questions. In the present study, we employed immunofluorescence and electron microscopy to study EP in a system that had previously been shown to model key elements of EP *in vivo*.^24^ Based on the duration of transit, we found that EP diverged into fast (generally < 10 minutes) and slow (generally > 60 minutes) forms. Neutrophils engaged in fast EP displayed ameboid motion, consistent with previous studies confirming a role for neutrophil cytoskeletal rearrangement in EP.^24^ By contrast, neutrophils engaged in slow EP assumed a rounded appearance, often near the cell nucleus and surrounded by the perinuclear ER.

These observations strongly suggest that fast and slow EP are distinct, although without specific tools to block either form, specific roles remain difficult to define. Tavassoli et al^26^ suggested that bone marrow cells might take a transmegakaryocytic route to enter the circulation in states of increased cell demand. Indeed, an increase of EP has been observed in rodents after LPS-induced peritonitis, a condition of enhanced hematopoiesis requiring rapid mobilization of bone marrow cells.^24,41^ Fast EP could be suited for such a role, although the advantage of passing through an MK instead of directly through the thin sinusoid wall is unclear. Perisinusoidal MKs could simply offer more surface area for transit, complementing direct egress in a process analogous to how transendothelial migration is thought to complement egress between endothelial cells in peripheral tissues.^42^ Alternately, since MKs participate actively in EP, it may be that they select particular neutrophils for such passage, for example on the basis of surface integrin activation.^24^ An additional possibility is that EP allows MKs to modulate neutrophil function during passage. We had previously demonstrated that EP allows neutrophils and MKs to engage in bidirectional transfer of membranes between cytoplasmic neutrophils and the DMS.^24^ Although our EM studies again showed neutrophils directly within the MK cytoplasm, it remains possible that not all neutrophils take this course. Some neutrophils could transit through MKs while remaining within emperisomes and thus topologically outside the MK, akin to transendothelial migration.^43^ If so, opportunities for membrane exchange could be limited, although soluble material or exosomes might still transfer.

A role for MKs in egress to the circulation would be consistent also with our observation that EP is performed preferentially by the most mature MKs. During megakaryopoiesis, MK precursors differentiate into fully mature MKs in the vascular bone marrow niche.^44^ Depending on their maturity, MKs interact differently with endothelial cells. While immature MKs mostly assume “planar contacts” with endothelial cells, mature MKs protrude podosome-like structures and eventually cytoplasmic processes through the endothelial cell layer into the bone marrow sinusoids.^45^ Thus, preferential conduct of EP by mature MKs restricts the process to cells in direct contact with the lumen of blood vessels. Alternately, enhanced EP by mature MKs could reflect their larger cell size and more extensive DMS, providing a larger cell surface for cell-cell contact and more space to accommodate neutrophils.

Intriguingly, approximately half of neutrophils remain within MKs for an extended period, a process we term slow EP. These cells exhibit a sessile morphology. Failing to observe any degraded neutrophils within MKs, we assume that most eventually exit, though since almost all slow EP events extended beyond our video observations we cannot confirm this assumption. De Pasquale et al^35^ proposed that EP might serve as a “sanctuary” for neutrophils in an unfavorable bone marrow environment, though why sanctuary should be necessary is unclear. Localization between the nucleus and the perinuclear ER raises additional novel possibilities, such as to “intercept” mRNA emerging from the nucleus or modulating ER function.^46,47^ Without direct evidence, all such possibilities remain purely speculative.

A central topological problem presented by EP is how neutrophils leave the emperisome to enter the MK cytoplasm.^21,24^ We approached this problem via EM of 45 EP events. The most common localization of neutrophils was within a clearly demarcated vesicle, termed the emperisome.^24^ In other instances, only a single membrane separated the cytoplasm of the neutrophil from that of the MK, consistent with intracytoplasmic residence (**Figure 4C_II_**). We observed intermediate steps in which part of the neutrophil remained in the emperisome while part exhibited contact with the cell cytoplasm (**Figure 4C_I_**). These images suggest penetration of the neutrophil through part of the vesicle wall, rather than for example wholesale resorption or disintegration of the emperisome membrane. Other images are more difficult to categorize definitively with respect to emperisome vs. cytoplasm, in particular where the neutrophil is surrounded by the DMS (**Figures 4B_I_–B_II_**). Earlier EM studies had described these cells as residing “loosely in the canalicular system”^48^, though further study will be require to understand the topological relationship of these cells with respect to the intra-vesicular, cytoplasmic, and extracellular compartments. The technical limitations of EM do not allow us to determine whether these morphological phases represent distinct neutrophil fates, restricted for example to either fast or slow EP, or instead sequential stages undertaken by many or most neutrophils during EP.

Our study has several important limitations. We studied murine EP, employing a useful but nevertheless *in vitro* system. How our findings translate to human and *in vivo* contexts remains unknown. While fast and slow EP appear distinct from each other, suggesting distinct functions, we could not here define these functions and cannot exclude the possibility that they fulfil similar roles. Further research is required to elucidate how neutrophils transition between different intermediate stages; whether EP modulates the behavior of neutrophils, MKs, or platelets; and whether MKs outside the bone marrow compartment, for example in the lung, also engage neutrophils and other cells via EP.

Despite these limitations, our studies provide the first evidence that MK EP is a heterogeneous process through which neutrophils may engage with MKs either for a short or long duration, interacting with intracellular structures including the emperisome, the DMS, the cytoplasm, the perinuclear ER, and potentially the nucleus itself. Preserved in all mammalian species studied, across millions of years of otherwise divergent evolution^21^, these observations suggest that EP will likely serve a range of roles to be defined through further investigations.

## Acknowledgements

F.Y.H. was funded by an MD fellowship from Boehringer Ingelheim Fonds. P.C. was funded by the Arthritis National Research Foundation and the Gilead Sciences Research Scholars Program in Rheumatology. F.A.R was funded by an MD fellowship from Boehringer Ingelheim Fonds. R.G-B. was funded by an MD fellowship from Boehringer Ingelheim Fonds, a physician-scientist development grant from the Medical Faculty Heidelberg, and a research grant from the German Society for Rheumatology. P.A.N. was supported by NIH/NIAMS awards R01AR065538, R01AR075906, R01AR073201, R21AR076630, P30AR070253, R56AR065538, NIH/NHLBI R21HL150575, a Lupus Research Alliance Target Identification in Lupus Grant, the Fundación Bechara, and the Arbuckle Family Fund for Arthritis Research.

## Authorship contributions

F.Y.H. designed and conducted experiments, analyzed data, and drafted the manuscript. P.C. designed and supervised experiments, analyzed data, and edited the manuscript. F.A.R. conducted experiments and analyzed data. R.G-B. analyzed data and edited the manuscript. P.A.N. conceptualized and supervised the study, analyzed data, and drafted the manuscript.

## Disclosure of conflicts of interest

The authors declare no competing interests.

## Supplementary videos

**Movie 1. Fast EP**. Mature MKs were stained with anti-CD41 AF488 (green) and co-incubated with fresh bone marrow cells stained with anti-Ly6G AF594 (red). DNA was stained with DRAQ5 (blue). Scale bar 10 µm. The neutrophil enters the MK on the right side, migrates through the MK cytoplasm and egresses on the opposite side within 10 minutes.

**Movie 2. Slow EP**. Mature MKs were stained with anti-CD41 AF488 (green) and co-incubated with fresh bone marrow cells stained with anti-Ly6G AF594 (red). DNA was stained with DRAQ5 (blue). Scale bar 10 µm. The neutrophil is already inside the MK at the beginning of the image acquisition and remains inside for at least 60 minutes showing no migration inside the MK.

**Movie 3. Distinct morpholology of fast and slow EP**. Mature MKs were stained with anti-CD41 AF488 (green) and co-incubated with fresh bone marrow cells stained with anti-Ly6G AF594 (red). DNA was stained with Hoechst 33342 (blue). Scale bar 10 µm. Two neutrophils undergoing slow EP assume a sessile state. A third neutrophil enters the MK after 31 minutes and extends dynamic membrane protrusions to propel itself through the MK cytoplasm, exiting within few minutes.

## Notes

### Competing Interest Statement

The authors have declared no competing interest.

## References

1. Ebaugh FGJ, Bird RM. The Normal Megakaryocyte Concentration in Aspirated Human Bone Marrow. Blood. 1951;6(1):75–80. doi:10.1182/blood.V6.1.75.75

2. Machlus KR, Italiano JE, Jr. The incredible journey: From megakaryocyte development to platelet formation. J Cell Biol. Jun 10 2013;201(6):785–96. doi:10.1083/jcb.201304054

3. Junt T, Schulze H, Chen Z, et al. Dynamic visualization of thrombopoiesis within bone marrow. Science. Sep 21 2007;317(5845):1767–70. doi:10.1126/science.1146304

4. Cunin P, Penke LR, Thon JN, et al. Megakaryocytes compensate for Kit insufficiency in murine arthritis. J Clin Invest. May 1 2017;127(5):1714–1724. doi:10.1172/jci84598

5. Cunin P, Nigrovic PA. Megakaryocytes as immune cells. J Leukoc Biol. Jun 2019;105(6):1111–1121. doi:10.1002/jlb.Mr0718-261rr

6. Beaulieu LM, Lin E, Morin KM, Tanriverdi K, Freedman JE. Regulatory effects of TLR2 on megakaryocytic cell function. Blood. Jun 2 2011;117(22):5963–74. doi:10.1182/blood-2010-09-304949

7. D’Atri LP, Etulain J, Rivadeneyra L, et al. Expression and functionality of Toll-like receptor 3 in the megakaryocytic lineage. J Thromb Haemost. May 2015;13(5):839–50. doi:10.1111/jth.12842

8. Shiraki R, Inoue N, Kawasaki S, et al. Expression of Toll-like receptors on human platelets. Thromb Res. 2004;113(6):379–85. doi:10.1016/j.thromres.2004.03.023

9. Undi RB, Sarvothaman S, Narasaiah K, Gutti U, Gutti RK. Toll-like receptor 2 signalling: Significance in megakaryocyte development through wnt signalling cross-talk and cytokine induction. Cytokine. Jul 2016;83:245–249. doi:10.1016/j.cyto.2016.05.007

10. Negrotto S, C JDG, Lapponi MJ, et al. Expression and functionality of type I interferon receptor in the megakaryocytic lineage. J Thromb Haemost. Dec 2011;9(12):2477–85. doi:10.1111/j.1538-7836.2011.04530.x

11. Beaulieu LM, Lin E, Mick E, et al. Interleukin 1 receptor 1 and interleukin 1beta regulate megakaryocyte maturation, platelet activation, and transcript profile during inflammation in mice and humans. Arterioscler Thromb Vasc Biol. Mar 2014;34(3):552–64. doi:10.1161/atvbaha.113.302700

12. Markovic B, Wu Z, Chesterman CN, Chong BH. Quantitation of soluble and membrane-bound Fc gamma RIIA (CD32A) mRNA in platelets and megakaryoblastic cell line (Meg-01). Br J Haematol. Sep 1995;91(1):37–42. doi:10.1111/j.1365-2141.1995.tb05241.x

13. Jiang S, Levine JD, Fu Y, et al. Cytokine production by primary bone marrow megakaryocytes. Blood. Dec 15 1994;84(12):4151–6.

14. Zhao M, Perry JM, Marshall H, et al. Megakaryocytes maintain homeostatic quiescence and promote post-injury regeneration of hematopoietic stem cells. Nat Med. Nov 2014;20(11):1321–6. doi:10.1038/nm.3706

15. Bruns I, Lucas D, Pinho S, et al. Megakaryocytes regulate hematopoietic stem cell quiescence through CXCL4 secretion. Nat Med. Nov 2014;20(11):1315–20. doi:10.1038/nm.3707

16. Finkielsztein A, Schlinker AC, Zhang L, Miller WM, Datta SK. Human megakaryocyte progenitors derived from hematopoietic stem cells of normal individuals are MHC class II-expressing professional APC that enhance Th17 and Th1/Th17 responses. Immunol Lett. Jan 2015;163(1):84–95. doi:10.1016/j.imlet.2014.11.013

17. Zufferey A, Speck ER, Machlus KR, et al. Mature murine megakaryocytes present antigen-MHC class I molecules to T cells and transfer them to platelets. Blood Adv. Sep 12 2017;1(20):1773–1785. doi:10.1182/bloodadvances.2017007021

18. Pariser DN, Hilt ZT, Ture SK, et al. Lung megakaryocytes are immune modulatory cells. J Clin Invest. Oct 20 2020;doi:10.1172/jci137377

19. Campbell RA, Schwertz H, Hottz ED, et al. Human megakaryocytes possess intrinsic antiviral immunity through regulated induction of IFITM3. Blood. May 9 2019;133(19):2013–2026. doi:10.1182/blood-2018-09-873984

20. Ren X, Wen W, Fan X, et al. COVID-19 immune features revealed by a large-scale single-cell transcriptome atlas. Cell. Apr 1 2021;184(7):1895-1913.e19. doi:10.1016/j.cell.2021.01.053

21. Cunin P, Nigrovic PA. Megakaryocyte emperipolesis: a new frontier in cell-in-cell interaction. Platelets. Aug 17 2020;31(6):700–706. doi:10.1080/09537104.2019.1693035

22. Humble JG, Jayne WH, Pulvertaft RJ. Biological interaction between lymphocytes and other cells. Br J Haematol. Jul 1956;2(3):283–94. doi:10.1111/j.1365-2141.1956.tb06700.x

23. Larsen TE. Emperipolesis of granular leukocytes within megakaryocytes in human hemopoietic bone marrow. Am J Clin Pathol. Apr 1970;53(4):485–9. doi:10.1093/ajcp/53.4.485

24. Cunin P, Bouslama R, Machlus KR, et al. Megakaryocyte emperipolesis mediates membrane transfer from intracytoplasmic neutrophils to platelets. Elife. May 1 2019;8 doi:10.7554/eLife.44031

25. Overholtzer M, Mailleux AA, Mouneimne G, et al. A nonapoptotic cell death process, entosis, that occurs by cell-in-cell invasion. Cell. Nov 30 2007;131(5):966–79. doi:10.1016/j.cell.2007.10.040

26. Tavassoli M. Modulation of megakaryocyte emperipolesis by phlebotomy: megakaryocytes as a component of marrow-blood barrier. Blood Cells. 1986;12(1):205–16.

27. Schmitt A, Jouault H, Guichard J, Wendling F, Drouin A, Cramer EM. Pathologic interaction between megakaryocytes and polymorphonuclear leukocytes in myelofibrosis. Blood. Aug 15 2000;96(4):1342–7.

28. Centurione L, Di Baldassarre A, Zingariello M, et al. Increased and pathologic emperipolesis of neutrophils within megakaryocytes associated with marrow fibrosis in GATA-1(low) mice. Blood. Dec 1 2004;104(12):3573–80. doi:10.1182/blood-2004-01-0193

29. Schmitt A, Drouin A, Massé JM, Guichard J, Shagraoui H, Cramer EM. Polymorphonuclear neutrophil and megakaryocyte mutual involvement in myelofibrosis pathogenesis. Leuk Lymphoma. Apr 2002;43(4):719–24. doi:10.1080/10428190290016809

30. Thiele J, Krech R, Choritz H, Georgii A. Emperipolesis--a peculiar feature of megakaryocytes as evaluated in chronic myeloproliferative diseases by morphometry and ultrastructure. Virchows Arch B Cell Pathol Incl Mol Pathol. 1984;46(3):253–63. doi:10.1007/bf02890314

31. Larocca LM, Heller PG, Podda G, et al. Megakaryocytic emperipolesis and platelet function abnormalities in five patients with gray platelet syndrome. Platelets. 2015;26(8):751–7. doi:10.3109/09537104.2014.994093

32. McGinnis E, Chipperfield KM. Striking emperipolesis in megakaryocytes of gray platelet syndrome. Blood. Jun 27 2019;133(26):2809. doi:10.1182/blood.2019000494

33. Di Buduo CA, Alberelli MA, Glembostky AC, et al. Abnormal proplatelet formation and emperipolesis in cultured human megakaryocytes from gray platelet syndrome patients. Sci Rep. Mar 18 2016;6:23213. doi:10.1038/srep23213

34. Sims MC, Mayer L, Collins JH, et al. Novel manifestations of immune dysregulation and granule defects in gray platelet syndrome. Blood. Oct 22 2020;136(17):1956–1967. doi:10.1182/blood.2019004776

35. de Pasquale A, Paterlini P, Quaglino D, Quaglino D. Emperipolesis of granulocytes within megakaryocytes. Br J Haematol. Jun 1985;60(2):384–6. doi:10.1111/j.1365-2141.1985.tb07429.x

36. Schulze H. Culture, Expansion, and Differentiation of Murine Megakaryocytes from Fetal Liver, Bone Marrow, and Spleen. Curr Protoc Immunol. Feb 2 2016;112:22f.6.1-22f.6.15. doi:10.1002/0471142735.im22f06s112

37. Eckly A, Heijnen H, Pertuy F, et al. Biogenesis of the demarcation membrane system (DMS) in megakaryocytes. Blood. Feb 6 2014;123(6):921–30. doi:10.1182/blood-2013-03-492330

38. Hartigan JA, Hartigan PM. The Dip Test of Unimodality. The Annals of Statistics. 1985;13(1):70–84.

39. Lee KP. Emperipolesis of hematopoietic cells within megakaryocytes in bone marrow of the rat. Vet Pathol. Nov 1989;26(6):473–8. doi:10.1177/030098588902600603

40. Bobik R, Dabrowski Z. Emperipolesis of marrow cells within megakaryocytes in the bone marrow of sublethally irradiated mice. Ann Hematol. Feb 1995;70(2):91–5. doi:10.1007/bf01834387

41. Tanaka M, Aze Y, Fujita T. Adhesion molecule LFA-1/ICAM-1 influences on LPS-induced megakaryocytic emperipolesis in the rat bone marrow. Vet Pathol. Sep 1997;34(5):463–6. doi:10.1177/030098589703400511

42. Filippi MD. Neutrophil transendothelial migration: updates and new perspectives. Blood. May 16 2019;133(20):2149–2158. doi:10.1182/blood-2018-12-844605

43. Muller WA. Mechanisms of leukocyte transendothelial migration. Annu Rev Pathol. 2011;6:323–44. doi:10.1146/annurev-pathol-011110-130224

44. Stegner D, vanEeuwijk JMM, Angay O, et al. Thrombopoiesis is spatially regulated by the bone marrow vasculature. Nat Commun. Jul 25 2017;8(1):127. doi:10.1038/s41467-017-00201-7

45. Eckly A, Scandola C, Oprescu A, et al. Megakaryocytes use in vivo podosome-like structures working collectively to penetrate the endothelial barrier of bone marrow sinusoids. J Thromb Haemost. Nov 2020;18(11):2987–3001. doi:10.1111/jth.15024

46. Köhler A, Hurt E. Exporting RNA from the nucleus to the cytoplasm. Nat Rev Mol Cell Biol. Oct 2007;8(10):761–73. doi:10.1038/nrm2255

47. Schwarz DS, Blower MD. The endoplasmic reticulum: structure, function and response to cellular signaling. Cell Mol Life Sci. Jan 2016;73(1):79–94. doi:10.1007/s00018-015-2052-6

48. Parmley RT, Kim TH, Austin RL, Alvarado CS, Ragab AH. Emperipolesis of neutrophils by dysmorphic megakaryocytes. Am J Hematol. Dec 1982;13(4):303–11. doi:10.1002/ajh.2830130405

